# Stochastic dynamics of genetic broadcasting networks

**DOI:** 10.1101/074401

**Authors:** Davit A. Potoyan, Peter G. Wolynes

## Abstract

The complex genetic programs of eukaryotic cells are often regulated by key transcription factors occupying or clearing out of a large number of genomic locations. Orchestrating the residence times of these factors is therefore important for the well organized functioning of a large network. The classic models of genetic switches sidestep this timing issue by assuming the binding of transcription factors to be governed entirely by thermodynamic protein-DNA affinities. Here we show that relying on passive thermodynamics and random release times can lead to a “time-scale crisis” of master genes that broadcast their signals to large number of binding sites. We demonstrate that this “time-scale crisis” can be resolved by actively regulating residence times through molecular stripping. We illustrate these ideas by studying the stochastic dynamics of the genetic network of the central eukaryotic master regulator *NFκB* which broadcasts its signals to many downstream genes that regulate immune response, apoptosis etc.

## INTRODUCTION

During development gene regulatory programs translate the information in the genome into phenotypes and after development these programs then micro-manage many functions within cells to ensure survival in a changing environment. At the molecular level, regulation involves an intricate web of interactions where the protein products of one set of genes bind to other genes and their products to modulate the production of biomolecules. The simplest element of a gene regulatory network is a genetic switch which can be thought of as a cellular control unit that can turn *ON* or *OFF* in response to external stimuli or signals from other genes [1]. The earliest models of genetic switches were formulated using purely thermodynamic models [2,3]. These models assumed that gene states are rapidly equilibrated 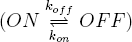 so that binding free energies, 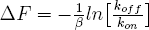, are the sole quantities controlling the systems level behavior of the network. These models, first introduced in studies on bacteria [4,5] are often used to explain gene expression patterns of higher organisms also in terms of protein-DNA interaction affinities. Thermodynamic models indeed have been fruitful in interpreting Chip-seq and binding microarray data and thereby have served as a conceptual link between molecular structure and gene expression [3,6]. In vivo, however, eukaryotic networks are generally far from equilibrium and the switching between states of a gene involves many elaborate kinetically controlled steps such as conformational changes of chromatin, assembly of various protein co-factors into larger transcription complexes, RNA polymerase attachment, etc. The apparently simple concept of a gene switching in response to equilibrium binding is therefore a high level idealization which may not be universally applicable.

Under time varying non-equilibrium conditions the *in vivo* activity of genes will be dictated not only by equilibrium binding but also by the residence time of transcription factors once they are bound to the DNA. Recent single cell and single molecule studies [7–9], show there are significant departures from the predictions of conventional thermodynamic models. Single-molecule chase assay experiments which directly measure the dissociation rates of transcription factors are physically inconsistent with the naive equilibrium models for the genetic switch [8]. A number of possibilities have been proposed to account for the departures from thermodynamic models [8,10]. Some have argued that energy consuming kinetic proofreading schemes could be employed by eukaryotic cells for attaining greater sensitivity [11] and specificity [12] of transcription factor binding relative to binding in equilibrium.

One intriguing mechanism that has been largely overlooked is that of an active regulation of the unbinding step through induced molecular stripping processes which remove transcription factors from their genomic binding sites. While some transcription factors may be spontaneously released from bound complexes the in vitro kinetic studies of the important transcription factor, *NFκB,* interacting with DNA and its inhibitor *IκB* have suggested that the regulation of *NFκB* in cells is likely to be kinetically controlled directly by the inhibitor [13, 14]. Molecular dynamics simulations have provided a detailed molecular level picture [15] of how *IκBα* strips the transcription factor off of DNA more rapidly than passive dissociation could. In addition, recent years single molecule level experiments have uncovered other cases of active regulation where protein-DNA exchange involves the formation of ternary complexes and consequent concentration dependent dissociation of proteins from DNA [7]. Facilitated dissociation has been observed in systems as diverse as the non-specifically bound architectural proteins [16, 17], metal sensing transcriptional regulators [9, 18], RNA polymerase [19] and even in ribosomal subunit switching [20].

Unlike the bacterial switches studied in the golden age of the molecular biology of bacteria, *NFκB* does not act as a simple switch turning on a single metabolic pathway but is involved in a very wide range of regulatory activities in eukaryotic cells [21]. To carry out these activities turning on the *NFκB* switch broadcasts a signal to many downstream genes. As we shall discuss in the paper, this broadcasting responsibility of *NFκB* creates a severe problem of timing if only passive dissociation of *NFκB* from its targets were possible. The conventional switch models that do not account for this active regulation of dissociation times encounter a “time scale crisis”. Owing to the large genomes and complex life styles of eukaryotes, *NFκB* like many other master switches has a huge number of target sites that initiate downstream functions [22, 23]. There are also myriads of non-functional sites where *NFκB* binds [24]. The *NFκB* activity coordinates a symphony of genes in response to complex environmental stimuli. Once the external environment returns to normal, however, not only is there no longer any need for further expression of these *NFκB* target genes but if they are not promptly turned off, deleterious actions may result. In contrast as we shall see by employing molecular stripping, the concentration of free and transcriptionally active *NFκB* will be promptly titrated back to zero by its inhibitor *IκB* once the stimulus is turned off. In many previous models of the *NFκB/IκB/DNA* circuitry *IκB* was thought to simply wait to encounter *NFκB* molecules that became unbound passively from the *DNA* in order to finally remove *NFκB* from the nucleus. We shall see that waiting for the NFkB to unbind from tens of thousands of sites in order to become available for the freely diffusing *IκB* simply takes too much time. Facing a changing environment, time becomes essential to the organism. The *IκB* induced direct stripping of *NFκB* from its genetic sites prevents this time scale crisis. Regulating residence times is a necessity for genes that broadcast signals to multiple targets.

### The NFκB/IκB/DNA broadcasting network

The transcription factors of the *NFκB* family are present in large quantities in eukaryotic cells (~ 10^5^ copies per cell). They activate ~ 5 · 10^2^ different genes [25, 26] in response to external stimuli. Due to its wide ranging influence over so many signaling activities *NFκB* is regarded as a master regulator switch which “broadcasts” signals to many target genes. Here we employ a stochastic model of the *NFκB* broadcasting network that includes the core inhibitory feedback loop of *IκB* along with the many DNA targets and decoy sites to which *NFκB* binds including most importantly the promoter for *IκB* itself (Table 1 and Supplementary Information). The target genes which are activated by the binding of *NFκB* to initiate signaling activities downstream are accompanied by a large majority of genomic sites that do not code for proteins and are likely decoys serving no known functions. An order of magnitude estimate for the number for decoys comes from genome-wide Chip-seq assays of binding [24] which have detected more than 2 · 10^4^ different DNA sites that bind *NFκB.* Our usage of the word “decoy” in the discussion below warrants some further explanation. Aside from the promoter of *IκB* all the other *NFκB* binding sites will be treated in the same way as effectively homogeneous binding sites obtained by dividing the accessible part of the genome into finite non overlapping stretches of DNA each of which binds a single *NFκB* molecule. Using the effective decoy concept allows us to average out sequence dependent effects. In the present contribution the effective binding sites are nearly identical entities with comparable residence times. The heterogeneity of binding/release properties of different sites is certainly an interesting issue (especially from the bioinformatic perspective) to which we plan to return in a future publication. The model network we study contains *D* = 2 · 10^4^ such effective decoys. The total number of *NFκB* molecules is 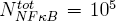. Both the numbers of decoys and the total number of *NFκB* molecules are essentially constant owing to the long cellular lifetime of *NFκB.* It should be noted that these effective decoys have larger capture rates and residence times for sliding along the DNA compared to the short consensus sequences of *NFκB* where binding has been studied *in vitro.* The parameters and reaction rates used in the model are shown in Table 1. The values for the rates of binding and unbinding steps are adopted from in vitro DNA binding experiments [13, 14] and genome wide microarray data [23]. The rest of the rates are coming from bulk kinetic experiments [27]. The binding ON rates are mostly diffusion limited and are set to *k*_*on*_ = *k*_*don*_ = *10 µMmin*^−1^. The OFF rates on the other hand show greater variation [23] and generally fall in the range between 10^−2^ and 1*min*^−1^. We vary the effective decoy OFF rates, *k*_*doff*_ in this range in order to obtain a complete survey of dynamic regimes that may be exhibited by this system or other analogous broadcasting networks. The OFF rate for the single *IκB* promoter site governs the period of oscillations and is set to its known value *k*_*off*_ ~ 0.1 *min*^−1^ generating 1–2 hour long oscillations consistent with single cell experiments [28]. The slowest time scale in the feedback loop corresponds to the degradation of *mRNA* and the fastest time scales correspond to the various binding events of *NFκB* to *IκB* and to the *DNA* sites. In our calculations dissociation from DNA bound sites can occur by passive unbinding with unimolecular kinetics 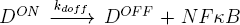 or by molecular stripping, i.e. active *IκB* concentration dependent dissociation with bimolecular kinetics 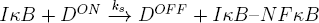

External stimulation of the network is modeled by setting the value of *α,* the rate of degradation of *IκB* in the bound *NFκB – IκB* complex. In real cells there are many sources of stimulation such as exposure to cytokines, free radicals, etc. All of these stimuli cause *IκB* to be marked for degradation thus freeing the cytoplasmic *NFκB* which then diffuses into the nucleus to begin the regulatory cycle. Under steady stimulation (setting *α = const*) the network settles into a self sustained oscillatory mode with periodic production and degradation of *IκB* molecules. The termination of stimulation corresponds to cessation of the degradation of *IκB* by setting *α* to 0. Figure 2 shows a sample of 50 trajectories of the time course of *NFκB-bound* decoys. These trajectories are initiated from a state where all of the decoys are bound to the *NFκB* with the remaining of *NFκB* molecules being part of the complex with *IκB.* We compare the cases with and without molecular stripping when the network is under constant external stimulation or transiently after that stimulation has been terminated. By looking at the switching when the stimuli have been terminated one immediately sees that even with our very conservative estimate of decoy residence times there is “time-scale crisis” for the conventional switch with no stripping. The time to turn off all the target genes exceeds by far the time scales of oscillation and gene expression (~ 120 – 150*min*). Meanwhile when there is molecular stripping the switching off of all the decoy and target genes happens in under ~ 20*min*. These time-scales are very much in harmony with what Fagerlund et al [30] have observed in the single cell studies probing the difference in nuclear clearance of *NFκB* for cells having wild type versus mutated *IκB* forms. Their experiments show that cells with wild type *IκB* rapidly and robustly clear *NFκB* from nucleus within a 20–30 min window but cells where *IκB* has been mutated in its PEST sequence which is crucial for molecular stripping [31] take ~ 100 – 150*min* in order to clear the *NFκB* from the nucleus. This observation is consistent with what one obtains from our model using reasonable values for decoy unbinding rates along with the rates in feedback loop that are inferred directly from bulk phase kinetic experiments. For the case of steady stimulation when only passive release occurs there are only highly stochastic oscillations with partial clearance of bound decoys. In contrast when there is molecular stripping, the system displays ultra-sensitive and highly regular pulsatile behavior in which decoys and targets are fully cleared in oscillations with well defined period. The differences between passive release switches and switches that employ molecular stripping for decoy clearance are explored in the next section.

**Fig. 1.**
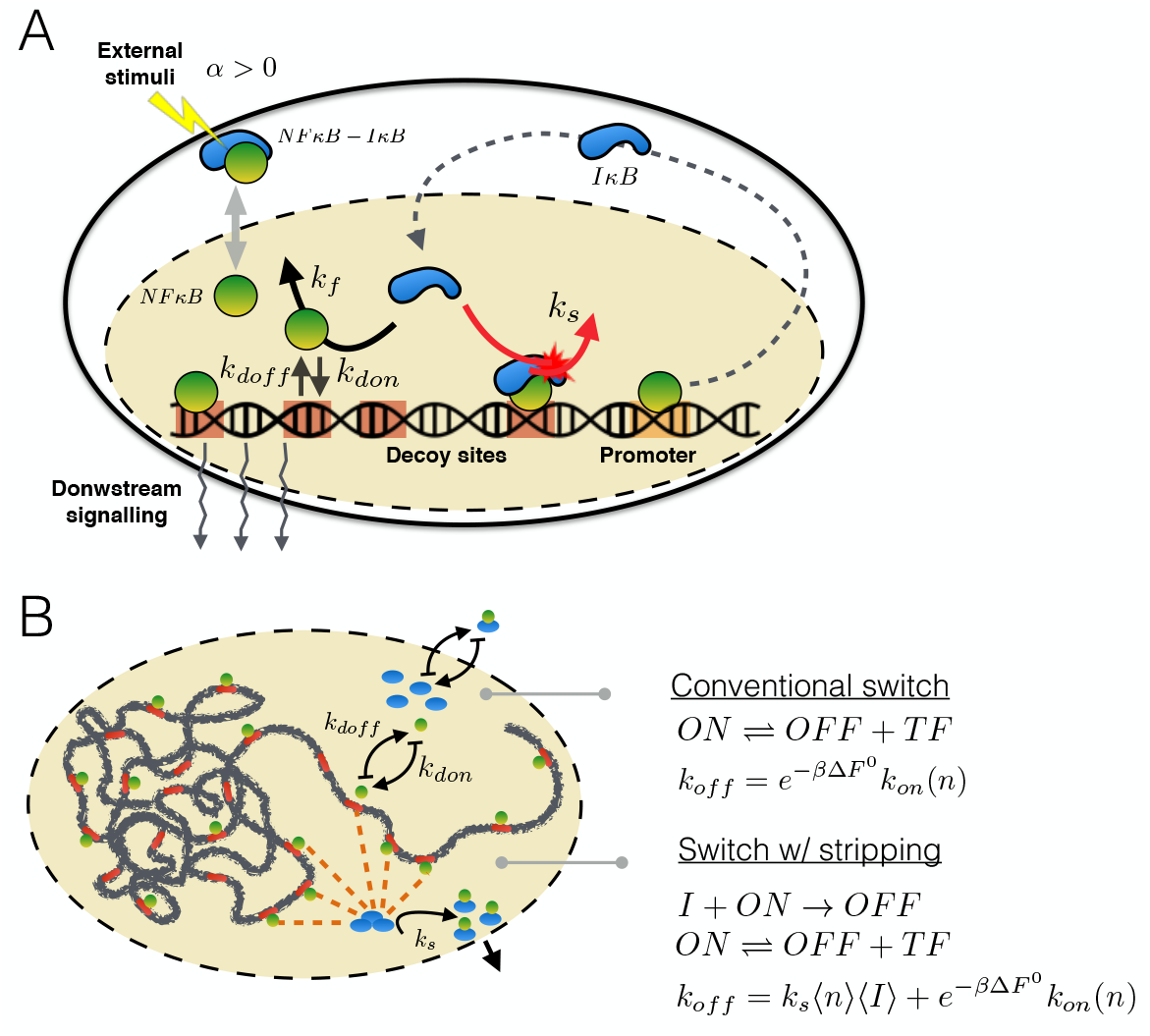
(A) Schematic picture of the broadcasting network of master regulator *NFκB.* Under external stimulation the *NFκB — IκB* is marked for degradation of *IκB* with the rate *α.* Once freed, *NFκB* goes inside nucleus and binds to myriad of DNA sites, including functional targets for downstream signaling, non-functional decoys and the promoter for *IκB* itself. Binding to the promoter site initiates the synthesis of *IκB.* The negative feedback loop of *IκB* is shown with dashed line with its inhibition of *NFκB* taking place either via direct binding to dissociated *NFκB* (solid black arrow) or via molecular stripping. (B) Shown are the mechanisms of 1) Conventional switch: where DNA unbinding rate takes place via passive dissociation and 2) Switch with stripping: where switching is being controlled kinetically via broadcasting signals of *IκB* stripping *NFκB* off of DNA sites directly in addition to passive dissociation step.

**Fig. 2.**
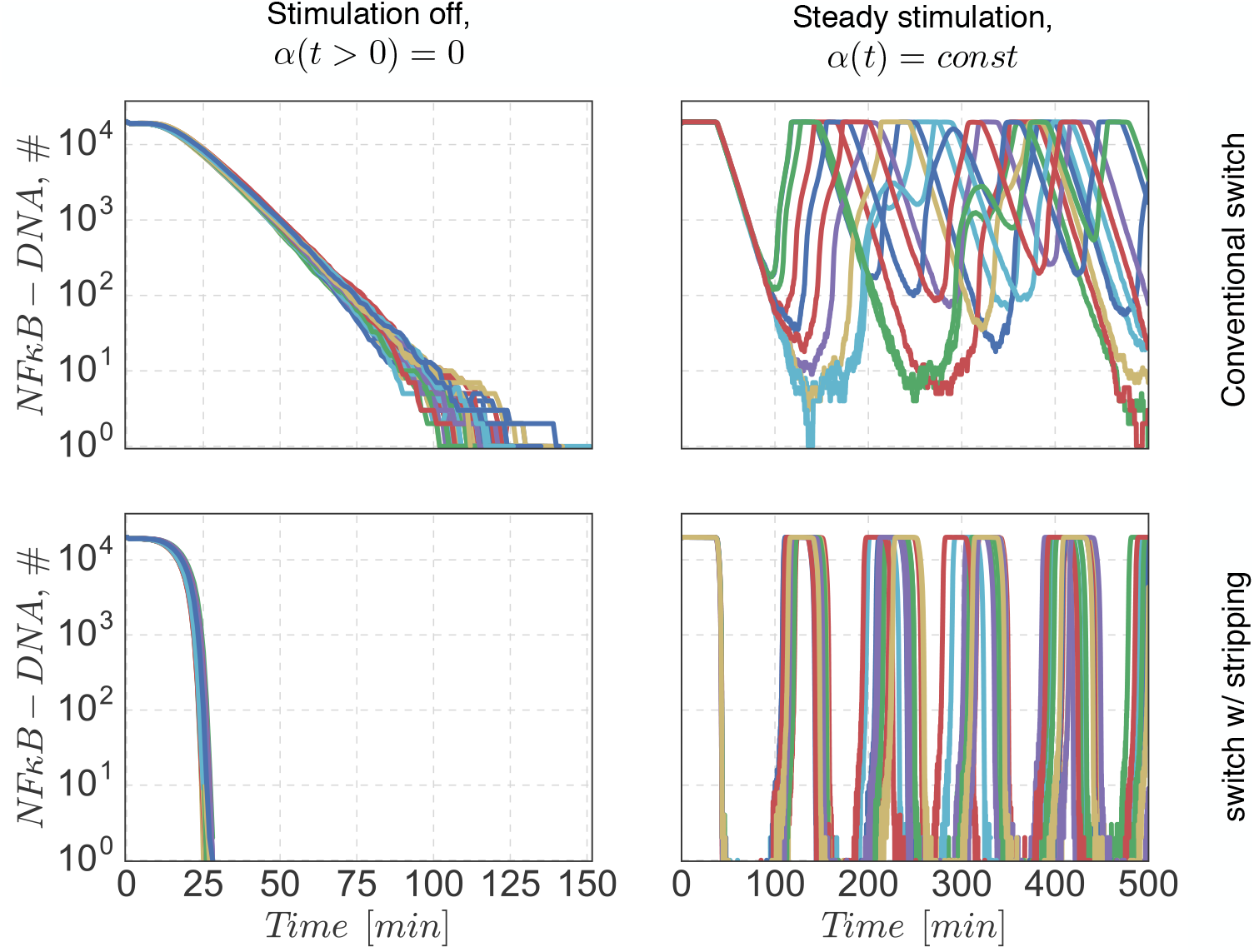
Shown are representative trajectories of networks functioning with conventional switches (first row) and switches with stripping (second row) under conditions of either terminated (*α* = 0) and continuous (*α >* 0) stimulation. Unbinding rates for all the bound decoys are *k*_*doff*_ = 0.1*min*^−1^.

**TABLE I.**
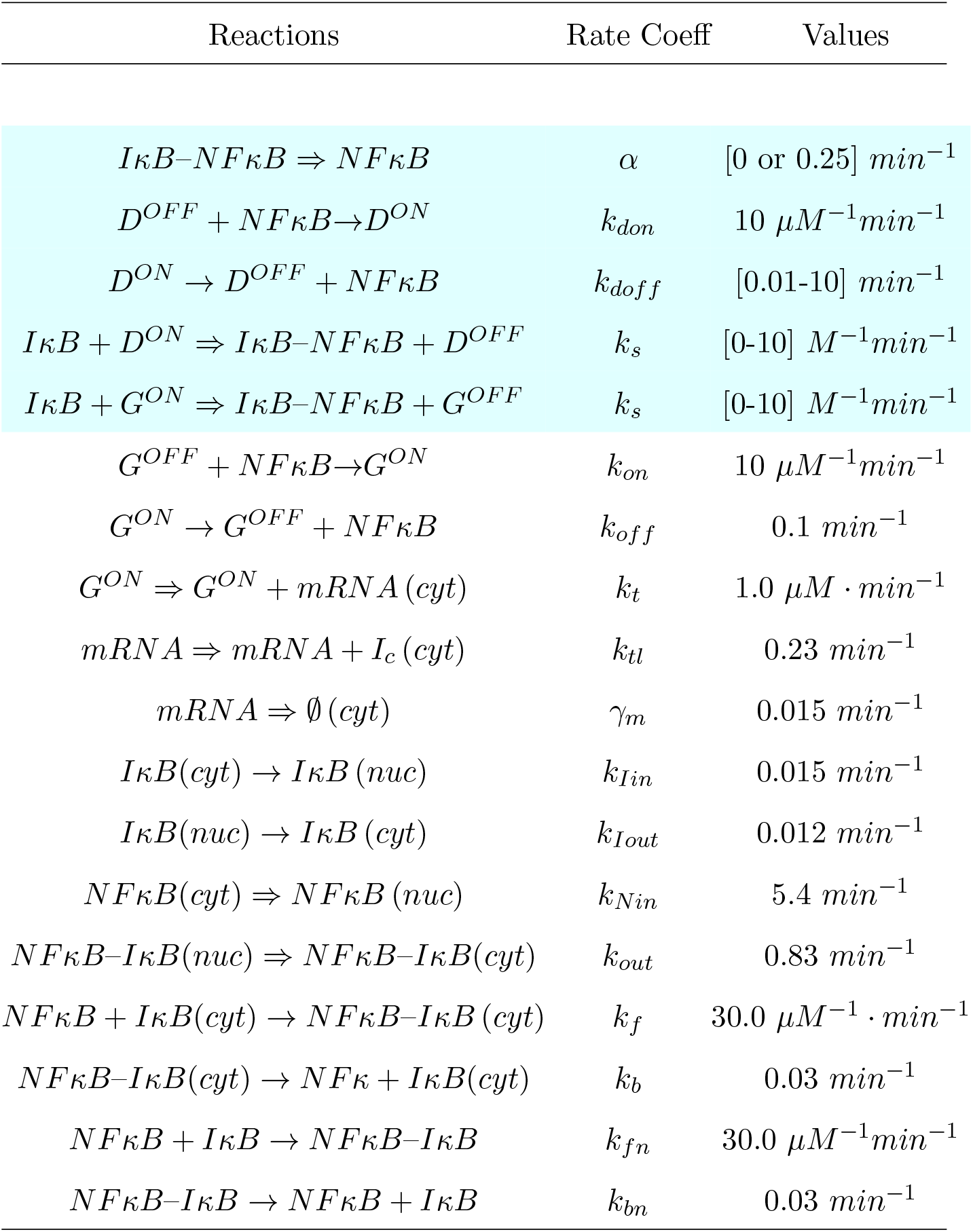
Reactions and rate coefficients for the *NFκB/IκB/DNA* master switch network. Dynamics of the network is simulated via a standard individual based kinetic Monte Carlo algorithm [29]. *G*^*ON*^, *G*^*OFF*^ denote *NFκB* bound and free state of *IκB* promoter. *D*^*ON*^, *D*^*OFF*^ denote *NFκB* bound and free states of decoys. Network has 1 *IκB* promoter and 2 · 10^4^ effective decoys. Fully irreversible reactions are indicated by ⇒ arrow. The kinetic parameters that are varied are highlighted in color. Reactions taking place in cytoplasm or transferring into cytoplasm from nucleus are indicated by (cyt) and (nuc) the rest are all happening in the nucleus.

| Reactions | Rate Coeff | Values |
| --- | --- | --- |
| $I\kappa B - NF\kappa B \Rightarrow NF\kappa B$ | $\alpha$ | $[0 \text{ or } 0.25] \text{ min}^{-1}$ |
| $D^{OFF} + NF\kappa B \rightarrow D^{ON}$ | $k_{don}$ | $10 \mu M^{-1} \text{ min}^{-1}$ |
| $D^{ON} \rightarrow D^{OFF} + NF\kappa B$ | $k_{doff}$ | $[0.01-10] \text{ min}^{-1}$ |
| $I\kappa B + D^{ON} \Rightarrow I\kappa B - NF\kappa B + D^{OFF}$ | $k_s$ | $[0-10] M^{-1} \text{ min}^{-1}$ |
| $I\kappa B + G^{ON} \Rightarrow I\kappa B - NF\kappa B + G^{OFF}$ | $k_s$ | $[0-10] M^{-1} \text{ min}^{-1}$ |
| $G^{OFF} + NF\kappa B \rightarrow G^{ON}$ | $k_{on}$ | $10 \mu M^{-1} \text{ min}^{-1}$ |
| $G^{ON} \rightarrow G^{OFF} + NF\kappa B$ | $k_{off}$ | $0.1 \text{ min}^{-1}$ |
| $G^{ON} \Rightarrow G^{ON} + mRNA(cyt)$ | $k_t$ | $1.0 \mu M \cdot \text{min}^{-1}$ |
| $mRNA \Rightarrow mRNA + I_c(cyt)$ | $k_{tl}$ | $0.23 \text{ min}^{-1}$ |
| $mRNA \Rightarrow \emptyset(cyt)$ | $\gamma_m$ | $0.015 \text{ min}^{-1}$ |
| $I\kappa B(cyt) \rightarrow I\kappa B(nuc)$ | $k_{Iin}$ | $0.015 \text{ min}^{-1}$ |
| $I\kappa B(nuc) \rightarrow I\kappa B(cyt)$ | $k_{Iout}$ | $0.012 \text{ min}^{-1}$ |
| $NF\kappa B(cyt) \Rightarrow NF\kappa B(nuc)$ | $k_{Nin}$ | $5.4 \text{ min}^{-1}$ |
| $NF\kappa B - I\kappa B(nuc) \Rightarrow NF\kappa B - I\kappa B(cyt)$ | $k_{out}$ | $0.83 \text{ min}^{-1}$ |
| $NF\kappa B + I\kappa B(cyt) \rightarrow NF\kappa B - I\kappa B(cyt)$ | $k_f$ | $30.0 \mu M^{-1} \cdot \text{min}^{-1}$ |
| $NF\kappa B - I\kappa B(cyt) \rightarrow NF\kappa B + I\kappa B(cyt)$ | $k_b$ | $0.03 \text{ min}^{-1}$ |
| $NF\kappa B + I\kappa B \rightarrow NF\kappa B - I\kappa B$ | $k_{fn}$ | $30.0 \mu M^{-1} \text{ min}^{-1}$ |
| $NF\kappa B - I\kappa B \rightarrow NF\kappa B + I\kappa B$ | $k_{bn}$ | $0.03 \text{ min}^{-1}$ |

### Active regulation of bound transcription factors resolves time-scale crisis in broadcasting networks

As we saw in the previous section without molecular stripping the sheer number of binding sites and the consequent long time to clear them pose serious problems for reliable operation of eukaryotic broadcasting switches. To explore the “time-scale crisis” more comprehensively we vary the dissociation rates of bound decoys over a wider range [10^−2^ – 10] going from normal dissociation rates to very fast rates. To quantify the robustness of the switching process we compute the mean first passage times for clearing all the decoys of their bound *NFκB* starting from a fully occupied decoy state once the stimulus has been terminated by setting *α*(*t* > 0) = 0 in the model. We computed over 10^4^ trajectories for each setup and measured the mean and coefficient of variation (*CV*) of the clearance time for different unbinding rates going from the regime of the pure conventional switch (*k*_*s*_ = 0 *M*^−^*min*^−1^) to moderate molecular stripping (*k*_*s*_ = 10 *M*^−1^*min*^−1^). Only for extremely fast intrinsic dissociation rates is the system able to function without molecular stripping (Fig 3). For more realistic values of the dissociation rates from effective decoys (~ 0.01 – 0.1) without stripping, a “time scale crisis” arises in which some target genes take dramatically more time to clear than the basic gene expression time scale and the time scale of oscillations. Once stripping is allowed the switch’s behavior is insensitive to orders of magnitude variation of molecular stripping rates (Fig 3A). Without stripping the need for the multiplicity of binding sites to unbind stochastically makes turning off the targets highly unpredictable with some genes lingering in their active states for extremely long times. This lingering effect is quantified by the coefficient of variation of clearance times as a function of unbinding rates shown in Fig 3B. From Fig 3 we see that the switch without molecular stripping gets noisier for sufficiently slow unbinding rates with the coefficient of variation diverging for slow dissociation rates of *NFκB* from decoys. This implies that different cells in the population would turn off their *NFκB* targets at random with some targets remaining ON for much longer times than others do. Molecular stripping, on the other hand curbs the noise consistently throughout the range of unbinding rates.

**Fig. 3.**
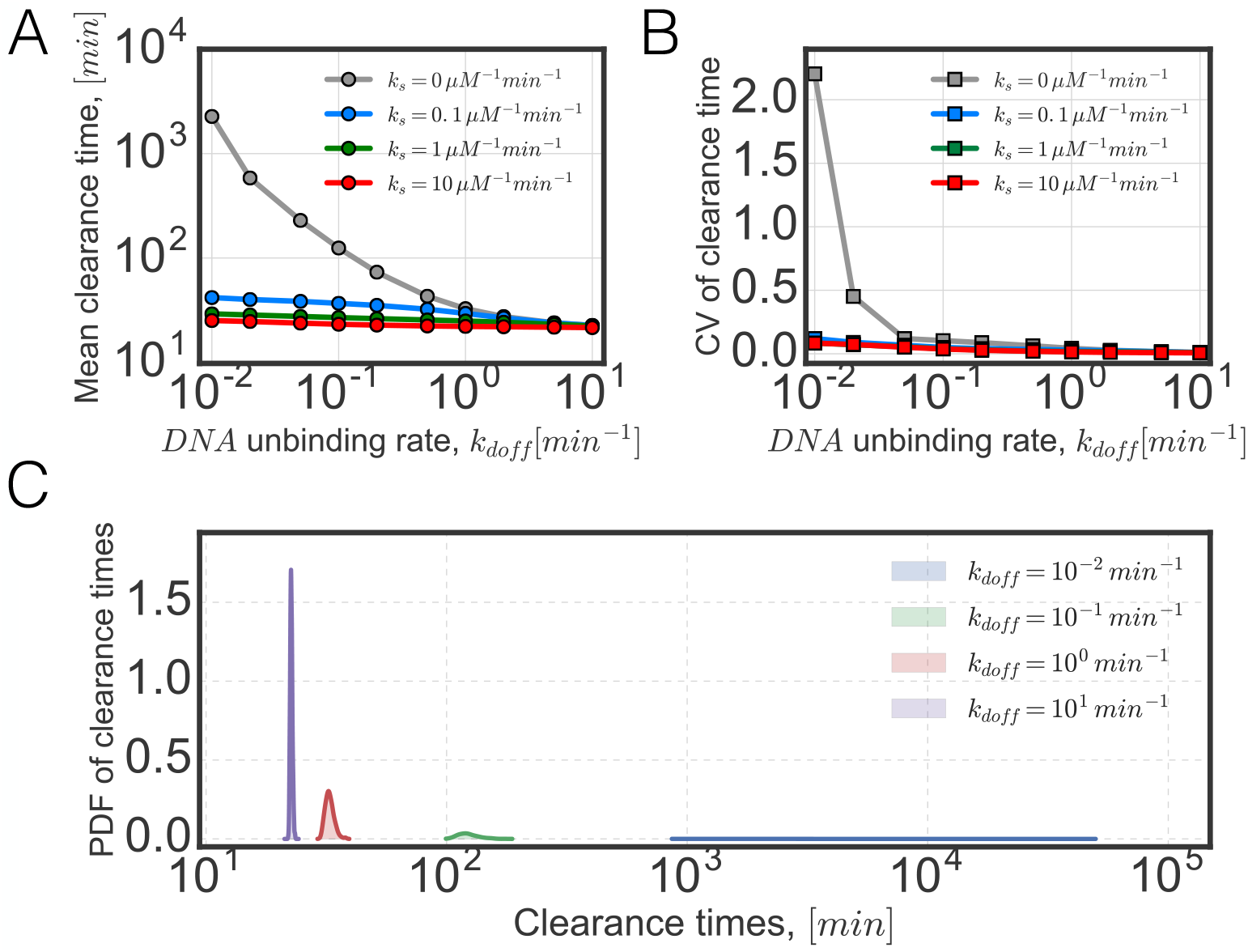
(A) Shown is the dependence of mean clearance time of occupied decoys, *NFκB – DNA* on unbinding rate for cases of no striping and stripping with rates *k*_*s*_. (B) Shown is the dependence of standard deviation of clearance times for cases of no striping and stripping with rates *k*_*s*_. (C) Shown are the probability distributions corresponding to different unbinding rates in the case of conventional switching, *k*_*s*_ = 0.

The divergence of the coefficient of variation for conventional switch has two origins, the large number of decoys and their slow dissociation rates. To see how these two factors contribute to the mean and coefficient of variation, we simulate the network with conventional switch while systematically varying the number of decoys and the dissociation rates (Fig 4). Within the biologically relevant parameter range the effect of this divergence is apparent for decoy numbers in the range of 10^3^ — 10^4^ and for dissociation rates from these decoys being ~ 10^−2^*min*^−1^ (Fig 4B). Surprisingly, when the dissociating rates are on the slow end of spectrum, of the order of ~ 10^−2^*min*^−1^ the mean clearance time already exceeds hours even for the moderately low number of decoys 10^2^ – 10^3^ (Fig 4A).

**Fig. 4.**
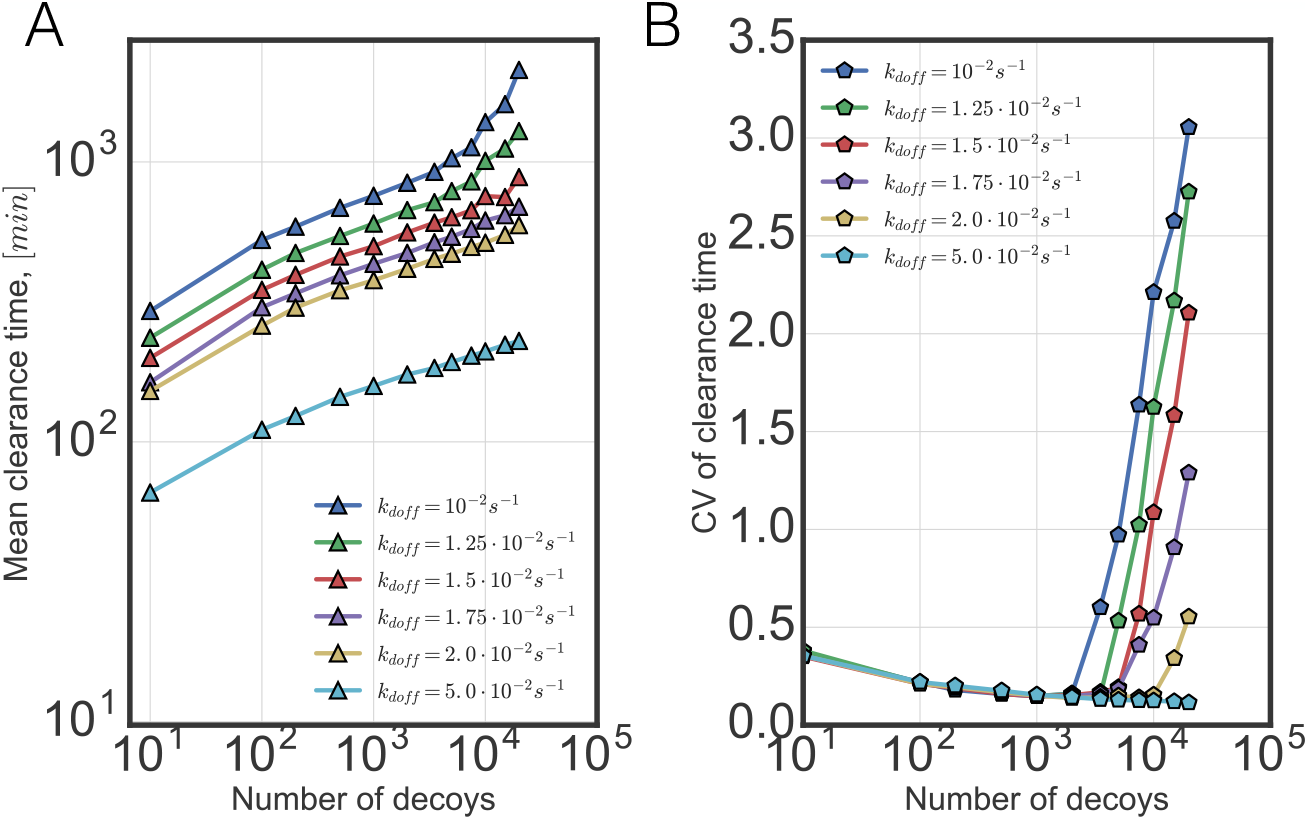
Shown are (A) mean and (B) coefficient of variation for the conventional switch as a function of numbers of decoys.

The stochastic nature of clearance is evident when we examine the stochastic dynamics of each individual decoy site (Fig 5A). Looking at the time trajectory of the occupancy of individual sites (quantified via probabilities of survival) reveals that some sites can remain occupied on time scales comparable to complete clearance without ever undergoing unbinding/rebinding (Supporting Information, Fig S1, S2). These “stragglers” would potentially pose problems for broadcasting networks without molecular stripping. When there is molecular stripping as we have seen from the Fig 3, clearance happens rapidly and reliably. These features are reflected in the rapidly decaying survival probabilities for individual DNA binding sites (Fig 5B). While the “stragglers” contribute greatly to the mean clearance time in the case of a conventional switch(Supporting Information, Fig S1), they are not the only source of variance. Rebinding events, where free *NFκB* binds back to the newly cleared sites not only slow down the complete clearance of bound *NFκB* but also make the clearance times highly unpredictable which is reflected in the divergence of the coefficient of variation seen in Fig 4. These rebinding events are purely stochastic in their origin, happening mostly at the later stages of clearance (Fig 5C) when the number of bound decoys is reduced to dozens and where the large number of unoccupied sites start to compete with the *IκB* for the *NFκB*(Supporting Information, Fig S1). Rebinding of *NFκB* to cleared decoy sites becomes more likely as the dissociation rates from the decoys becomes comparable to the rate of dissociation of *NFκB* from its complex with the *IκB* (Fig 5C). When there is molecular stripping, the rebinding events simply do not matter as the rate of clearance is too rapid for rebinding to influence clearance (Fig 5D).

**Fig. 5.**
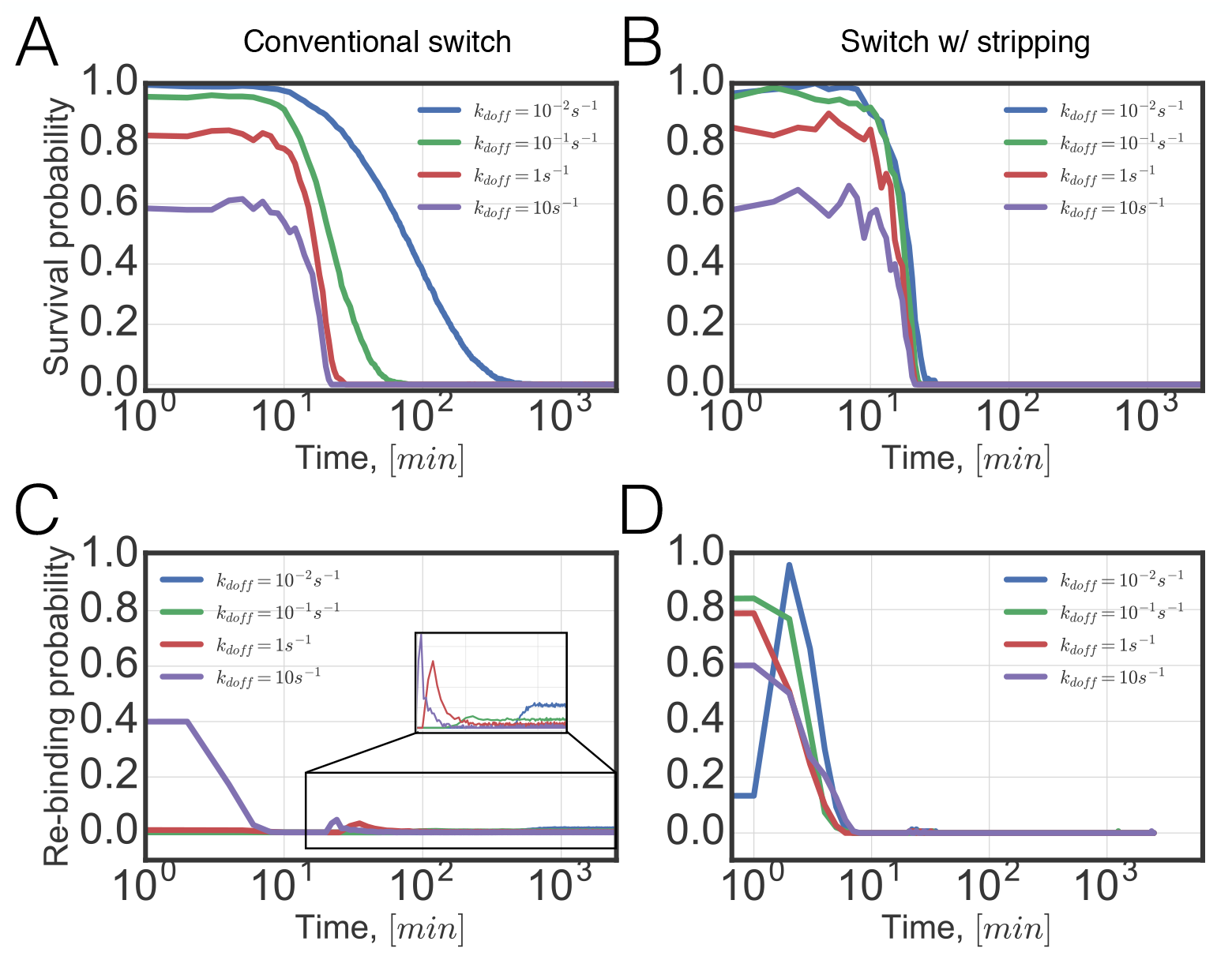
Shown are the survival probabilities of single bound decoy sites as a function of time for the networks (A) with no stripping and (B) with stripping. Rebinding probabilities are computed as a function of time for the cases (C) with no stripping and (D) with stripping. Different curves correspond to different dissociation rates of the *NFκB* bound decoys.

Now we turn to investigating how molecular stripping changes the stochastic behavior of the switch when the broadcasting system displays self-sustained pulsatile behavior. Keeping *α*(*t >* 0) = 0.25*min*^−1^ constant in the simulation leads to oscillations of the nuclear *NFκB* with a period of ~ 2*hr* (Supplementary Information, Fig S4) consistent with commonly adopted experimental setup [28]. In the present study, however we are interested in the consequences for the residence times of *NFκB* at genomic binding sites. Therefore we examine the oscillations of the occupation of decoy states as a function of unbinding rates. The mean fraction of the time that the sites are completely cleared (Fig 6A) provides a good way to quantify how long the genes are transcriptionally silent on average under the external stimulation. Different patterns of pulses of *NFκB* are thought to activate different patterns of genes for downstream signaling [32]. Experiments suggest the temporal dynamics of *NFκB* has information that can be utilized for decision making by the cell. For the conventional switch without any molecular stripping the mean fraction of completely cleared times is essentially zero for a broad range of dissociation rates as there are always some *NFκB* molecules bound to the decoy sites at all times. Only in the fast dissociation limit do the oscillations become clearly pulsatile alternating between completely cleared and occupied states. With molecular stripping, due to the fast turnover of the *NFκB* the oscillations appear to be ultra sensitive over a wide range of values of molecular stripping rate. A curious situation occurs in the fast dissociation limit where the passive switch slightly edges out the switch gene itself which has the slow molecular stripping rate of *k*_*s*_ ~ 0.1 *M*^−1^*min*^−1^. This happens because the stripping of the promoter site for *IκB* reduces the overall rate of stripping of the remaining decoy sites beyond that of passive dissociation. That this is the appropriate explanation is verified by simulating what happens when one selectively turns off the stripping only at the promoter site. This change makes the mean wait times for all *k*_*s*_ > 0 even longer (See Supplementary Figure S3). One may imagine similar strategies used by the cell where the signals are being broadcast to many targets with a few selected ones gaining special protection by modification of chromatin structure. To quantify the variation in the bimodality of the signals of *NFκB — DNA* pulses we compute the complexity of the steady state probability distributions at various time points in the oscillation via Shannon entropy *S* = – Σ_*n*_ *p*(*n*)*log*_2_ *p*(*n*) where *p*(*n*) is the probability density of *NFκB — DNA* established after many pulses (Fig 6B). Comparing Shannon entropies of pulsatile stochastic trajectories for broadcasting networks we see that network without stripping generates more complex distributions which reflects the more unpredictable (or “surprising”) nature of oscillations. When there is molecular stripping, the distributions generated by pulses are more predictable as is reflected by low values of Shannon entropy. How pulsatile signals get corrupted by noise can be quantified by computing the relative entropy between stochastic and deterministic pulses, 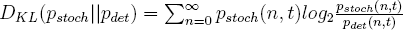 for time windows containing roughly one pulse (Fig 6C). As expected more irregular oscillations generated in the absence of molecular stripping deviate from pure deterministic pulses the most. For biologically relevant values of the stripping rates the stochastic pulses stay much closer to their respective deterministic ones. The auto-correlation of the pulses is another quantity that shows how rapidly noise randomizes the relative phases of different oscillators (Fig 6C). Again, networks with molecular stripping hold the phases correlated over much longer times than happens in the absence of striping where it takes at most 2–3 pulses for phases to become completely uncorrelated.

**Fig. 6.**
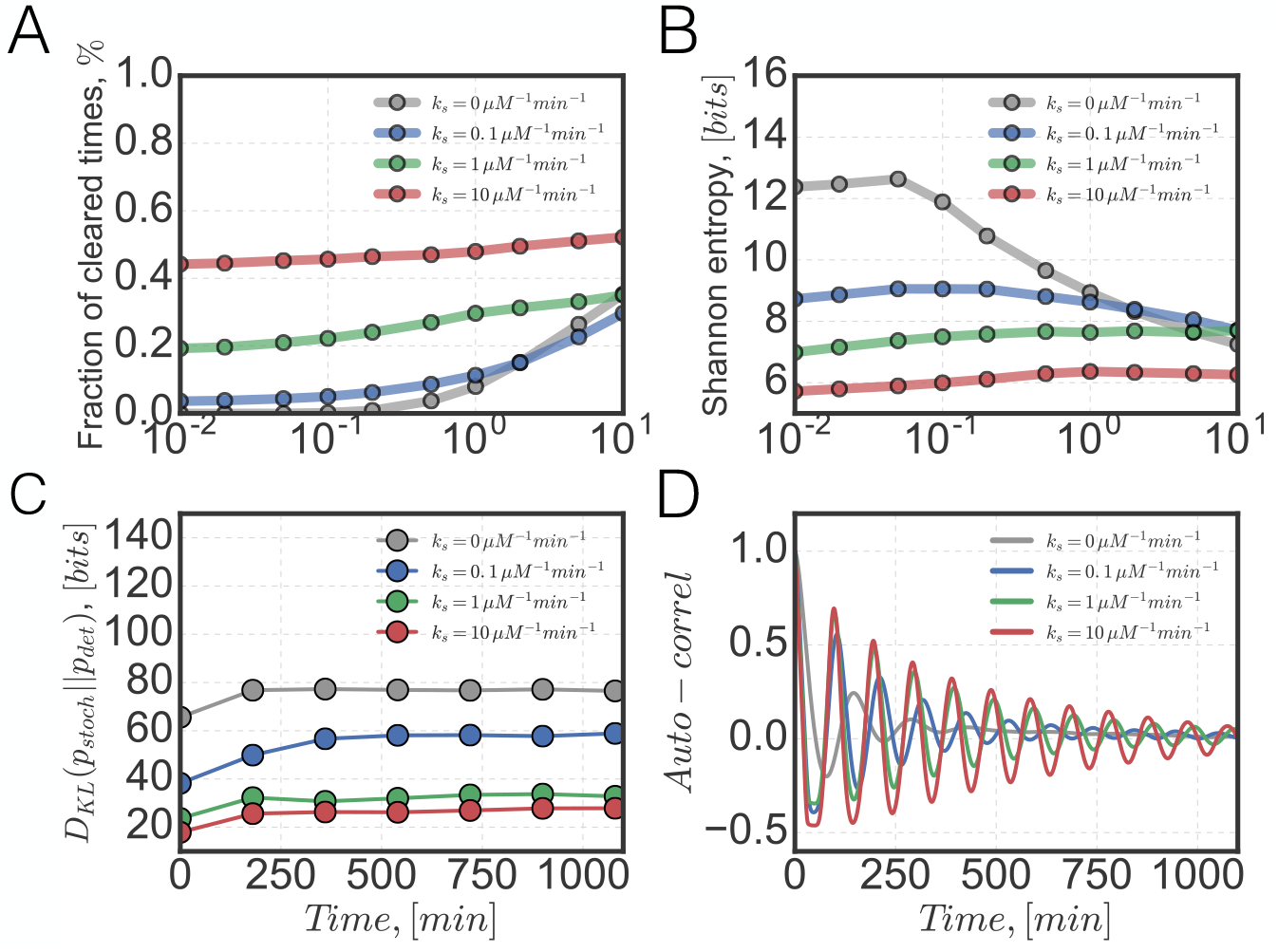
(A) Dependence of mean fraction of cleared times on decoy unbinding rate computed for networks different stripping rates *k*_*s*_. (B) Shannon’s entropy of steady state distribution of pulsatile signals as a function of decoy unbinding rate computed for different stripping rates *k*_*s*_. (C) Evolution of relative entropy (Kullback-Leibler) between stochastic and deterministic pulses computed in time windows containing roughly one pulse for the unbinding rate of *k*_*doff*_ = 0.1 *min*^−1^. (D) Temporal autocorrelation function of DNA bound *NFκB* molecules for the unbinding rate of *k*_*doff*_ = 0.1 *min*^−1^.

### The non-equilibrium nature of switches in a broadcasting system

Molecular stripping modifies the often adopted equilibrium picture of a gene switch by adding to the otherwise reversible step of transcription factor binding/unbinding a microscopically irreversible step. One should note however that in reality molecular stripping just like any other chemical reaction must have a finite backward rate, which in the case of stripping and a few other reactions in our network is thought to be vanishingly small. The network as a whole is driven outside of equilibrium by continuous stimulation from the outside. Therefore there is no question that both with and without stripping the *NFκB* regulatory network operates under highly non-equilibrium conditions where molecules are constantly being pumped into and degraded out of the system thereby guaranteeing the stability of steady oscillatory or homeostatic states. Quantifying the extent of irreversibility of genetic networks is not trivial partly because of the great complexity of real biological networks with many steps that could potentially contribute to the feedback cycle. One measure one can employ is the rate of dissipation which is also the rate of entropy production by the network caused by the explicitly detailed steps (i.e. not including the dissipative costs of protein synthesis which are clearly required too). The dissipation rate is computed by calculating averages of stochastic entropy production over the ensemble of trajectories 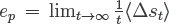. The stochastic entropy production is computed by taking ratios of probabilities of forward and backward transitions involving changes of species for each elementary stochastic event in the network: 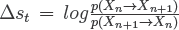. If the network were in equilibrium the stochastic entropy production would be exactly equal to zero. Entropy production turns out to be nonzero either with stripping or without it. Since the thermodynamic affinities of DNA sites are commonly used for thinking about gene regulation, we would like to quantify how informative such quantities are under the more general non-equilibrium conditions. To do this we first compute the entropy production by varying the unbinding rates while also keeping the equilibrium dissociation constant fixed (Fig 7). As expected regardless of the values of equilibrium constant the network is driven far from equilibrium. The networks with molecular stripping are more dissipative than those without stripping when the unbinding rates are slow. We see that the irreversible step of decoy clearance via molecular stripping gives rise to ultra-sensitive oscillations but does so at a cost of higher rates of dissipation. For the faster unbinding rates the many steps of *NFκB* capture and release take over causing the intrinsic entropy production by network without stripping to exceed that of the one with stripping.

**Fig. 7.**
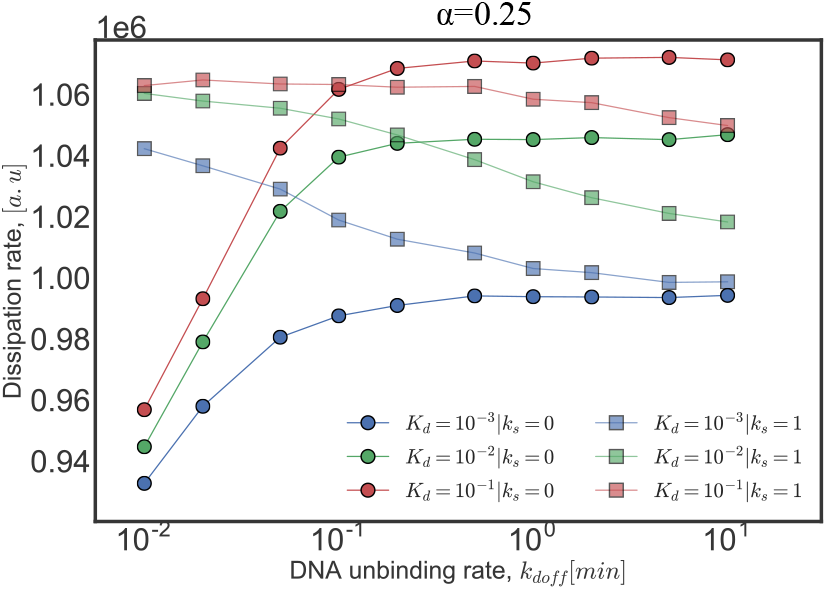
Rate of entropy production in the oscillatory steady state as a function of dissociation rate of bound decoys, *k*_*doff*_. Dissociation rates *kdoff* are varied by keeping the thermodynamic affinities fixed *K*_*d*_ = *k*_*doff*_/*k*_*don*_ = *const.* Different curves correspond to networks with and without stripping and to different values of dissociating constant.

When there is no stripping and when binding and spontaneous unbinding are fast, the equilibrium affinities are strongly correlated with the mean clearance time, with higher affinities corresponding to slower clearance and vice versa (Fig 8A). When binding and spontaneous unbinding are slow however, the mean clearance times weakly depend on the equilibrium affinities but are instead dictated by the dissociation rates. In a sense when there is molecular stripping the concept of affinity loses its meaning altogether. The means and the variances of the clearance times become independent of the equilibrium affinities (Fig 8A–B). At the steady state stripping causes the mass action ratios to be governed only by the binding ON rates since the effective OFF rates from all the decoys become equal(Fig 8C–D). These rates are very often diffusion limited. The equilibrium dissociation constant is clearly not a very reliable measure of the “strength” of binding for dynamical situations.

**Fig. 8.**
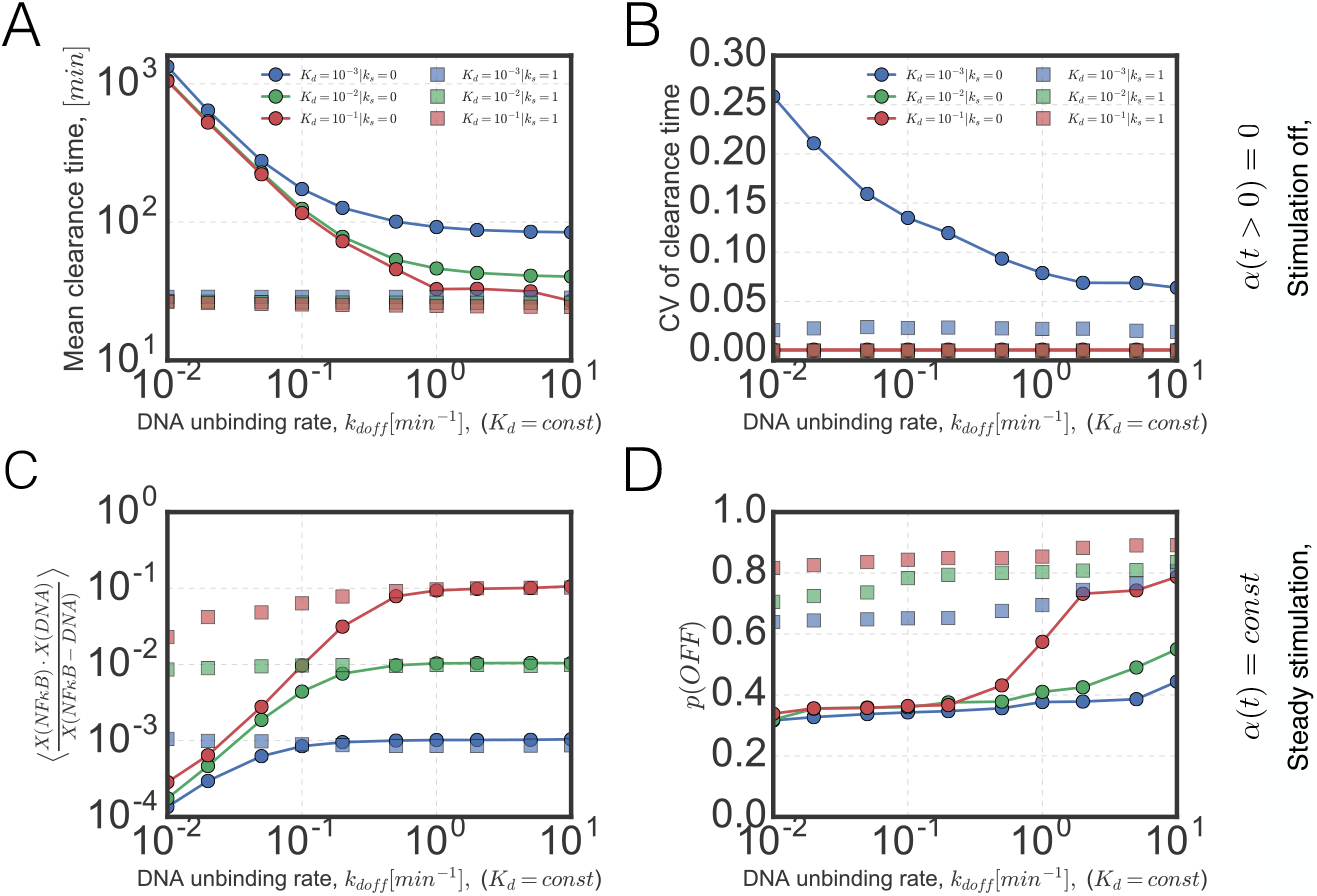
Shown are the values of (A) Mean clearance time and (B) coefficient of variation of the network with terminated stimuli as a function of dissociating rates, *k*_*doff*_ with fixed thermodynamic affinities *K*_*d*_ = *k*_*doff*_ /*k*_*don*_ = *const.* (C) Mass action ratio of decoy-*NFκB* quasi equilibrium, with the corresponding macroscopic dissociation as a function of dissociation rates *k*_*doff*_. (D) Probability of unoccupied *IκB* promoter.

## Conclusion

In this work we have studied the stochastic dynamics of the broadcasting genetic network centered on *NFκB.* Physiological, genomic and bioinformatic studies have shown that *NFκB* is a master regulator which binds to huge number of DNA sites having coding, downstream regulatory and non-functional consequences. Single cell experiments have shown that under steady stimulation the network exhibits self sustained oscillations but that once the external stimuli are terminated the network rapidly switches into a non-oscillatory homeostatic state. Here we have put forward a new conceptual framework for understanding the mechanism of regulation of broadcasting networks at the systems level. The large number of targets for the master regulator necessitates going beyond conventional models of gene regulation which were inspired by studies on simple bacterial systems with few targets. We show that passive dissociation of transaction factors leads to a ‘time-scale crisis’ for broadcasting signals to targets if the targets and decoys are too numerous. In the case of the *NFκB* network, molecular stripping by *IκB* solves the ‘time-scale crisis’ and leads to fast resposnes and ultra-sensitive oscillations with well defined periods. Our results are in harmony with recent *in vivo* single cell experiments where the stripping process has been perturbed specifically.

## References

[1] M. Ptashne, A genetic switch: phage lambda revisited, Vol. 3 (Cold Spring Harbor Laboratory Press Cold Spring Harbor, NY:, 2004).

[2] G. K. Ackers, A. D. Johnson, and M. A. Shea, Proc Natl Acad Sci 79, 1129 (1982).

[3] L. Bintu, N. E. Buchler, H. G. Garcia, U. Gerland, T. Hwa, J. Kondev, and R. Phillips, Current opinion in genetics & development 15, 116 (2005).

[4] W. Gilbert and B. Müller-Hill, Proc Natl Acad Sci 56, 1891 (1966).

[5] F. Jacob and J. Monod, J Mol Biol 3, 318 (1961).

[6] G. D. Stormo and Y. Zhao, Nature Reviews Genetics 11, 751 (2010).

[7] T. Ha, Cell 154, 723 (2013).

[8] P. Hammar, M. Walldén, D. Fange, F. Persson, O. Baltekin, G. Ullman, P. Leroy, and J. Elf, Nature genetics 46, 405 (2014).

[9] T.-Y. Chen, A. G. Santiago, W. Jung, L. Krzemiński, F. Yang, D. J. Martell, J. D. Helmann, and P. Chen, Nature communications 6 (2015).

[10] A. Coulon, C. C. Chow, R. H. Singer, and D. R. Larson, Nat Rev Gen 14, 572 (2013).

[11] J. Estrada, F. Wong, A. DePace, and J. Gunawardena, Cell 166, 234 (2016).

[12] S. A. Cepeda-Humerez, G. Rieckh, and G. Tkačik, Phys Rev Lett 115, 248101 (2015).

[13] S. Bergqvist, V. Alverdi, B. Mengel, A. Hoffmann, G. Ghosh, and E. A. Komives, Proc Natl Acad Sci 106, 19328 (2009).

[14] V. Alverdi, B. Hetrick, S. Joseph, and E. A. Komives, Proc Natl Acad Sci 111, 225 (2014).

[15] D. A. Potoyan, W. Zheng, E. A. Komives, and P. G. Wolynes, Proc Natl Acad Sci USA 113, 110 (2016).

[16] J. S. Graham, R. C. Johnson, and J. F. Marko, Nucleic acids research 39, 2249 (2011).

[17] M. J. McCauley, E. M. Rueter, I. Rouzina, L. J. Maher, and M. C. Williams, Nucleic acids research 41, 167 (2013).

[18] C. P. Joshi, D. Panda, D. J. Martell, N. M. Andoy, T.-Y. Chen, A. Gaballa, J. D. Helmann, and P. Chen, Proc Natl Acad Sci 109, 15121 (2012).

[19] J. J. Loparo, A. W. Kulczyk, C. C. Richardson, and A. M. van Oijen, Proc Natl Acad Sci USA 108, 3584 (2011).

[20] D. D. MacDougall and R. L. Gonzalez, Journal of molecular biology 427, 1801 (2015).

[21] H. L. Pahl, Oncogene 18 (1999).

[22] R. Martone, G. Euskirchen, P. Bertone, S. Hartman, T. E. Royce, N. M. Luscombe, J. L. Rinn, F. K. Nelson, P. Miller, M. Gerstein, S. Weissman, and M. Snyder, Proc Natl Acad Sci USA 100, 12247 (2003).

[23] T. Siggers, A. B. Chang, A. Teixeira, D. Wong, K. J. Williams, B. Ahmed, J. Ragoussis, I. A. Udalova, S. T. Smale, and M. L. Bulyk, Nature immunology 13, 95 (2012).

[24] B. Zhao, L. A. Barrera, I. Ersing, B. Willox, S. C. Schmidt, H. Greenfeld, H. Zhou, S. B. Mollo, T. T. Shi, K. Takasaki, et al., Cell reports 8, 1595 (2014).

[25] A. Antonaki, C. Demetriades, A. Polyzos, A. Banos, G. Vatsellas, M. D. Lavigne, E. Apostolou, E. Mantouvalou, D. Papadopoulou, G. Mosialos, et al, J Biol Chem 286, 38768 (2011).

[26] S. C. Gupta, C. Sundaram, S. Reuter, and B. B. Aggarwal, Biochim Biophys Acta 1799, 775 (2010).

[27] A. Hoffmann, A. Levchenko, M. L. Scott, and D. Baltimore, Science 298, 1241 (2002).

[28] D. Nelson, A. Ihekwaba, M. Elliott, J. Johnson, C. Gibney, B. Foreman, G. Nelson, V. See, C. Horton, and D. Spiller, Science 306, 704 (2004).

[29] D. T. Gillespie, J Phys Chem 81, 2340 (1977).

[30] R. Fagerlund, M. Behar, K. T. Fortmann, Y. E. Lin, J. D. Vargas, and A. Hoffmann, J Roy Soc Int 12, 20150262 (2015).

[31] D. A. Potoyan, W. Zheng, D. U. Ferreiro, P. G. Wolynes, and E. A. Komives, J Phys Chem B 120, 8532 (2016).

[32] S. Zambrano, I. De Toma, A. Piffer, M. E. Bianchi, and A. Agresti, Elife 5, e09100 (2016).

